# Patterns of neuronal activation following ethanol-induced social facilitation and social inhibition in adolescent cFos-LacZ male and female rats

**DOI:** 10.1101/2024.03.06.583793

**Authors:** Trevor T. Towner, Devon T. Applegate, Harper J. Coleman, Elena I. Varlinskaya, David F. Werner

## Abstract

Motives related to the enhancement of the positive effects of alcohol on social activity within sexes are strongly associated with alcohol use disorder and are a major contributor to adolescent alcohol use and heavy drinking. This is particularly concerning given that heightened vulnerability of the developing adolescent brain. Despite this linkage, it is unknown how adolescent non-intoxicated social behavior relates to alcohol’s effects on social responding, and how the social brain network differs in response within individuals that are socially facilitated or inhibited by alcohol. Sex effects for social facilitation and inhibition during adolescence are conserved in rodents in high and low drinkers, respectively. In the current study we used cFos-LacZ transgenic rats to evaluate behavior and related neural activity in male and female subjects that differed in their social facilitatory or social inhibitory response to ethanol. Subjects were assessed using social interaction on postnatal days 34, 36 and 38 after a 0, 0.5 and 0.75 g/kg ethanol challenge, respectively, with brain tissue being evaluated following the final social interaction. Subjects were binned into those that were socially facilitated or inhibited by ethanol using a tertile split within each sex. Results indicate that both males and females facilitated by ethanol display lower social activity in the absence of ethanol compared to socially inhibited subjects. Analyses of neural activity revealed that females exhibited differences in 54% of examined socially relevant brain regions of interest (ROIs) compared to only 8% in males, with neural activity in females socially inhibited by ethanol generally being lower than facilitated subjects. Analysis of socially relevant ROI neural activity to social behavior differed for select brain regions as a function of sex, with the prefrontal cortex and nucleus accumbens being negatively correlated in males, but positively correlated in females. Females displayed additional positive correlations in other ROIs, and sex differences were noted across the rostro-caudal claustrum axis. Importantly, neural activity largely did not correlate with locomotor activity. Functional network construction of social brain regions revealed further sex dissociable effects, with 90% interconnectivity in males socially inhibited by ethanol compared to 38% of facilitated subjects, whereas interconnectivity in females inhibited by ethanol was 10% compared to nearly 60% in facilitated subjects. However, hub analyses converged on similar brain regions in males and females, with the nucleus accumbens being a hub region in socially inhibited subjects, whereas the central amygdala was disconnected in facilitated subjects. Taken together, these findings support unified brain regions that contribute to social facilitation or inhibition from ethanol despite prominent sex differences in the social brain network.

## Introduction

Alcohol use is typically initiated during early adolescence (Faden 2006), with alcohol use becoming normative in the United States by about 15 years of age (Masten et al. 2009). Initiation of drinking before this age (Dawson et al., 2008), especially drinking to intoxication in early adolescence (Morean et al. 2014) is associated with adverse outcomes, including the development of alcohol use disorder (AUD). It is not surprising that the developing brain is extremely sensitive to different insults including alcohol use given substantial neurodevelopmental changes that occur during adolescence (Lees et al. 2020; Spear 2018). According to the Substance Abuse and Mental Health Services Administration’s (SAMHSA) National Survey on Drug Use and Health in 2019, 2.3 million adolescents aged 12 to 17 initiated alcohol use, which corresponds to approximately 6,200 adolescents initiating alcohol use each day (SAMHSA 2020). Among this age group, 414,000 adolescents were diagnosed with AUD, with more adolescent females than males (251,000 versus 163,000, respectively) meeting criteria for AUD (SAMHSA 2020).

These epidemiological data suggest sex differences in sensitivity to detrimental effects of alcohol during adolescence, with adolescent females being more sensitive than their male counterparts. Indeed, adolescent girls have been shown to be more susceptible to adverse consequences of binge drinking than adolescent boys, reporting greater anxiety and depression than adolescent males following a recent binge episode (Bekman et al. 2013). While many adolescents use alcohol, not all of them develop alcohol-related problems, suggesting a substantial role of individual differences in alcohol sensitivity. There is some evidence associating increased risk for AUD with accentuated sensitivity to the positive and/or stimulatory effects of alcohol as well as attenuated sensitivity to the negative and sedative effects (King et al. 2011; Quinn and Fromme 2011). Together, these findings suggest that individual differences as well as sex differences play a substantial role in adolescent vulnerability to alcohol.

During the adolescent developmental period, interactions with peers become extremely important, with adolescents spending more time with peers than younger or older individuals (Lam et al. 2014). Adolescents are also more sensitive to peer acceptance and rejection than children or adults (Andrews et al. 2021), with sensitivity to social reward also being heightened during adolescence (Foulkes and Blakemore 2016). Therefore, it is not surprising that adolescents drink predominantly in social situations (Anderson and Brown 2010; Terry-McElrath et al. 2017) and often report social (drinking to have a good time at social gathering) and enhancement (drinking to have fun) motives for drinking (Kuntsche et al. 2014; Smit et al. 2022). Not only motives, but also alcohol expectancies play a substantial role in adolescent alcohol use. Indeed, an expectancy for social facilitation from drinking is an important predictor of heavy drinking in adolescence, with young individuals believing that alcohol will make them more confident and relaxed in a social setting (Mackintosh et al. 2006). Therefore, alcohol-associated social facilitation plays a substantial role in adolescent alcohol use.

Adolescent rodents also demonstrate sex differences in sensitivity to ethanol, with adolescent females, for instance, being more sensitive to aversive properties of ethanol than adolescent males under social test circumstances (Morales et al. 2014). Likewise, ethanol-induced social facilitation is not restricted to human adolescents, but also evident in adolescent rats (Varlinskaya and Spear 2015). We have shown that systemic administration of low dose ethanol increased social interaction in adolescent rats that were tested on postnatal day (P) 28 and P35 (Varlinskaya and Spear 2002; 2006), with no ethanol-induced social facilitation evident in older adolescent rats tested on P42 (Varlinskaya and Spear 2006). While sensitivity to socially facilitating effect of ethanol declines with age, responding to socially inhibiting and anxiogenic effect of ethanol shows an ontogenetic increase (Varlinskaya and Spear 2015). Our previous research has revealed very limited sex differences in adolescent sensitivity to ethanol-induced social facilitation in between-subject designs when the social effects of acute ethanol were assessed relative to saline-injected controls (Varlinskaya and Spear 2002; 2006). Using a rat model of adolescence and within-subject design that allowed us to assess effects of ethanol relative to saline in the same animal, we revealed individual differences in sensitivity to the ethanol dose of 0.75 g/kg in adolescent males and females, with high drinkers demonstrating social facilitation and low drinkers showing social inhibition (Varlinskaya et al. 2015). These findings suggest individual differences in sensitivity to ethanol-induced social facilitation or social inhibition following acute challenge with 0.75 g/kg ethanol. It is still unknown, however, whether similar neural mechanisms contribute to ethanol-induced social facilitation (or social inhibition) in males and females.

It is likely that brain regions implicated in modulation of social interaction are differentially affected by ethanol in socially facilitated versus socially inhibited adolescents. The social brain circuitry is comprised of multiple brain regions including, but not limited to, the nucleus accumbens, extended amygdala, and lateral septum (Bickart et al. 2014; Gangopadhyay et al. 2021; Kas et al. 2014). Furthermore, specific neuronal ensembles within these regions may contribute to ethanol-induced changes in social responding, with these ensembles potentially being different in ethanol-induced social facilitation and social inhibition. To expand our understanding of individual differences in responding to the social effects of ethanol, it is important to characterize the neuronal activation patterns associated with social facilitation and social inhibition induced by a certain ethanol dose as well as to assess whether these patterns differ in adolescent males and females.

In the present study, we aimed to identify neuronal activation patterns associated with ethanol-induced social facilitation and ethanol-induced social inhibition following an acute challenge with 0.75 g/kg ethanol in male and female adolescent cFos-LacZ rats (Figure 1A,B). Since these transgenic rats express β-galactosidase (β-gal) under the control of the *cfos* promoter, cFos and β-gal proteins are co-expressed in neurons strongly activated by a certain stimulus and not in the surrounding neurons that are inactive or weakly activated (Cruz et al. 2014). Therefore, β-gal expression is used as a proxy for cFos (see Fig.1C). Expression of β-gal was assessed in the following regions of interest (ROIs): lateral orbitofrontal cortex (lOFC), ventral orbitofrontal cortex (vOFC), prelimbic cortex (PrL), infralimbic cortex (IL), anterior claustrum (aCL), posterior claustrum (pCL), nucleus accumbens core (NAcC), nucleus accumbens shell (NAcSh), dorsomedial striatum (DMS), dorsolateral striatum (DLS), lateral septum (LS), central amygdala (CeA), and basolateral amygdala (BLA). We hypothesized that brain regions associated predominantly with social approach behavior and social reward would have greater neuronal activation in adolescent rats that display ethanol-induced social facilitation, whereas brain regions more implicated in social anxiety-like behavior would have elevated neuronal activation in adolescents socially inhibited by ethanol.

**Figure 1.**
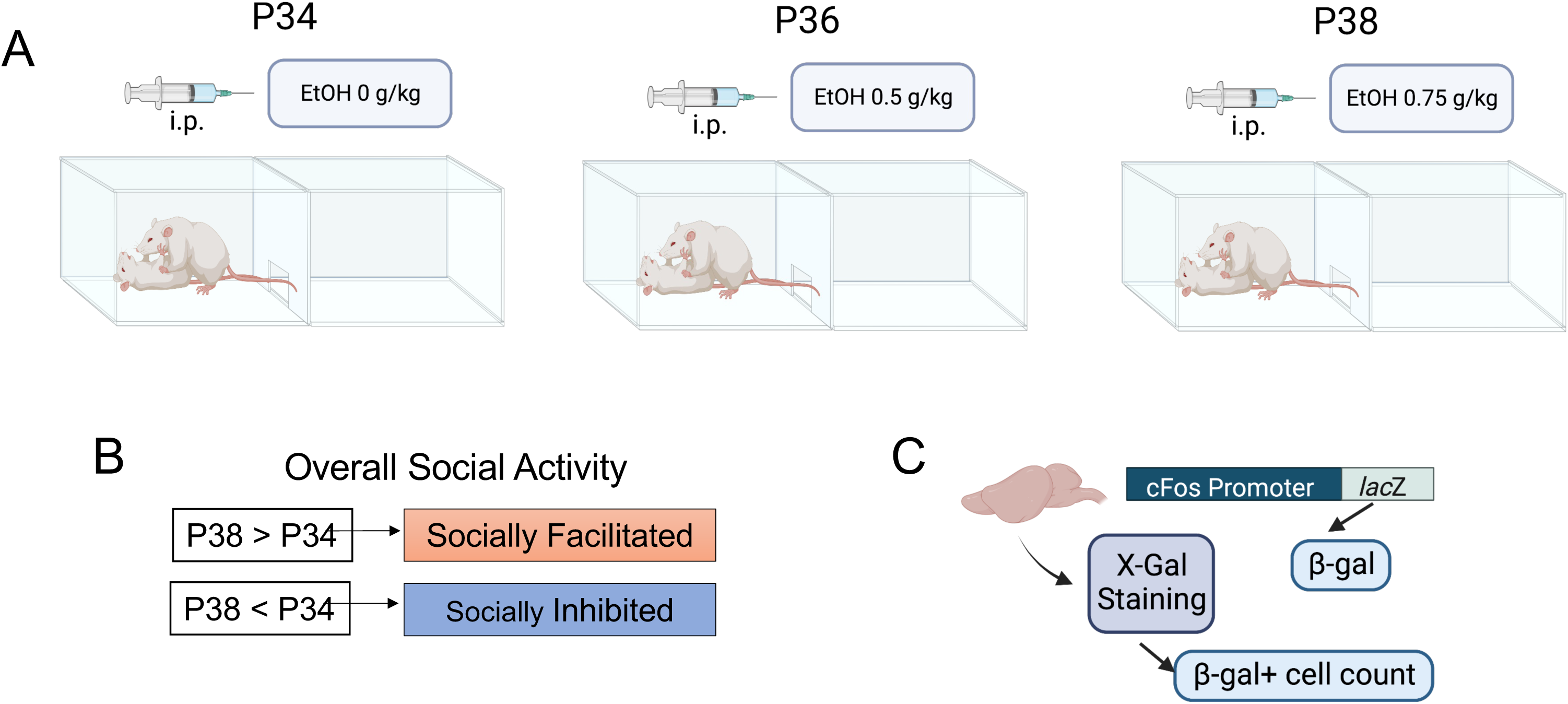
Ethanol-induced social facilitation and social inhibition in cFos-LacZ adolescent rats. (A) Experimental timeline; (B) assessments of social facilitation and social inhibition; (C) c-Fos-LacZ transgenic model of measuring β-galactosidase (β-gal) expression as a proxy for cFos.

## General Methods

### Subjects

Adolescent male and female cFos-LacZ transgenic rats on a Sprague Dawley background bred and reared in our colony at Binghamton University were used. Transgenic breeders were derived from the transgenic line originally developed by Dr. Curran while at Roche Institute of Molecular Biology and maintained by the Children’s Hospital of Philadelphia. For breeding, cFos-LacZ males were backcrossed with outbred Sprague-Dawley female breeders. On postnatal day (P)1, litters were culled to 8 – 10 pups, with equal sex ratios when possible. On P21, pups were weaned and pair-housed with same sex and age peers. Between P20 and P24 tissue was collected by ear punch for LacZ genotyping (TransnetYX, Cordova, TN). Only animals with the c*fos-lacZ transgene (LacZ positive)* were used for subsequent experiments. All animals were kept in a temperature controlled (22°C) vivarium maintained on a 12:12 hour light/dark cycle beginning at 0700 hours. Animals were given *ad libitum* access to food and tap water. All experimental procedures and maintenance of rats were in accordance with the National Institutes of Health guidelines for animal care using protocols approved by Binghamton University Institutional Animal Care and Use Committee.

### Procedure

Male and female experimental subjects were socially tested following an intraperitoneal (IP) injection of 0 g/kg ethanol (physiological saline vehicle) on P34, followed by an IP administration of 0.5 g/kg ethanol on P36, and an 0.75 g/kg ethanol on P38 (see Figure 1A for timeline). Ethanol was administered as 12.6% (v/v) solution. All injections were given immediately prior to placing of experimental subjects into the social interaction apparatus. On the final test day (P38), experimental animals remained with the social partner for a total of 60 minutes prior to brain tissue collection.

### Modified Social Interaction (SI) test

As in our previous studies (Towner et al. 2022; Varlinskaya et al. 2020), social behavior was assessed using the modified SI test. Immediately after an IP injection, experimental subjects were placed into a testing apparatus (30 x 20 x 20 cm) consisting of two equally sized compartments separated by a wall with an aperture allowing for free movement between the compartments. Pine shavings were placed at the bottom of the apparatus, and 6% hydrogen peroxide was used for cleaning the apparatus after each testing. Testing took place between 1100 and 1400 h in the presence of a white noise generator and under dim lighting (10 – 15 lux). Rats were placed into the testing apparatus alone for a 30-minute habituation period prior to introduction of a non-familiar, non-manipulated same-sex social partner that weighed about 10 g less than the experimental animal. On P34 and P36, experimental subjects and their partners were left together for 10 minutes, and this 10-minute SI test was videorecorded. On P38, each experimental subject was exposed to a novel social partner for 60 minutes, with only the first 10 minutes recorded. All social partners were wild type Sprague Dawley rats.

Overall social and locomotor activity were scored and analyzed. The sum of frequencies of social investigation (sniffing of any part of the body of the partner), contact behavior (crawling under and over the partner and social grooming), and play fighting (playful nape attack, chasing, and pinning) characterized overall social activity. The total number of crossovers (movements between the two compartments) performed by each experimental subject was used as an index of general locomotor activity during the SI test.

### Individual Differences in Ethanol Sensitivity: Tertile Split

To assess individual differences in sensitivity to ethanol effects on overall social activity, we calculated the percent change of social behavior from saline to the highest ethanol dose (0.75 g/kg). We then divided animals into ethanol social responding phenotype groups using a tertile split: socially facilitated (top third) by ethanol animals demonstrated substantial increases in overall social activity following injection of the 0.75 g/kg ethanol dose relative to their own values of overall social activity following saline injection, rats in the middle tertile did not demonstrate ethanol-induced changes, whereas in socially inhibited (bottom third) adolescent males and females, the dose of 0.75 g/kg produced decreases of overall social activity relative to saline. Only socially facilitated (n = 10 females; n = 8 males) and socially inhibited (n = 10 females; n = 8 males) animals were used in the present study (see Figure 1B).

### Brain Tissue Sectioning and Enzymatic Histology (X-Gal)

On the final test day (P38), experimental animals remained with the social partners for a total of 60 minutes and were then decapitated. Brains were rapidly removed, drop-fixed in 4% paraformaldehyde for 90 minutes, and then placed in 30% sucrose in phosphate buffered saline (PBS) until equilibrated, followed by flash freezing in methyl butane. Using a cryostat (Leica Biosystems, Buffalo Grove, IL) 30μm slices were obtained and placed into an antifreeze solution (30% ethylene glycol and 30% glycerol and 0.5M PBS) and stored at -20°C. X-gal histochemistry and visualization were conducted as we have done previously (Towner et al., 2022). In brief, coronal slices were initially agitated in fix buffer (0.1M phosphate buffer supplemented with 5mM EGTA, 2mM magnesium chloride and 0.2% glutaraldehyde) for 15 minutes followed by two 5-minute washes in wash buffer (0.1M phosphate buffer supplemented with 2mM magnesium chloride). Tissue slices were then placed in staining buffer (0.1M phosphate buffer supplemented with 2mM magnesium chloride, 5mM potassium ferrocyanide, 5mM potassium ferricyanide and 1mg/ml of X-Gal in 0.1M PBS) and incubated overnight at 37°C with gentile agitation. Tissue slices were then washed twice in wash buffer before being mounted onto slides and air-dried overnight. Slices were counterstained with eosin and dehydrated via ascending ethanol concentrations prior to submerging in xylenes and coverslipping with permount (Electron Microscopy Sciences, Hatfield, PA).

### Regions of Interest (ROIs) and Cell Counting

Thirteen ROIs were examined for expression of β-gal, namely the lOFC, vOFC, PrL, IL, aCL, pCL, NAcC, NAcSh, DMS, DLS, LS, CeA, and BLA (see Table 1 for coordinates. Stained slices were imaged with an Olympus Research Slide Scanner VS200 (Olympus, PA, US) at a 10x magnification. ROIs were identified from coronal images and cropped for image analysis with Olympus ASW software. Images were transferred to HALO image analysis platform (Indica Labs, NM, US), and β-gal positive (β-gal+) cells were automatically identified based on the expression of indigo blue staining and on soma size and roundness. The automated identification of β-gal+ cells was verified using hand-counted images with over 95% reliability between observation methods. All β-gal+ cell counts were converted to cell number per mm^2^.

**Table 1.** Regions of Interest (ROIs). Coordinates based on Paxinos and Watson, 2007

| Regions of Interest | Grid Size | Coordinates from Bregma |
| --- | --- | --- |
| Lateral orbitofrontal cortex (IOFC) | 0.4mm<sup>2</sup> | 5.16 – 3.00mm |
| Ventral orbitofrontal cortex (vOFC) | 0.4mm<sup>2</sup> | 5.16 – 3.00mm |
| Prelimbic cortex (PrL) | 0.4mm<sup>2</sup> | 4.68 – 2.52mm |
| Infralimbic cortex (IL) | 0.4mm<sup>2</sup> | 3.72 – 2.52mm |
| Anterior claustrum (aCL) | 0.3mm x 0.2mm | 3.72 – 2.52mm |
| Posterior claustrum (pCL) | 0.2mm x 0.3mm | 2.28 – 1.20mm |
| Nucleus accumbens core (NAcC) | 0.4mm<sup>2</sup> | 2.76 – 1.08mm |
| Nucleus accumbens shell (NAcSh) | 0.4mm<sup>2</sup> | 2.76 – 0.96mm |
| Dorsomedial striatum (DMS) | 1.0mm<sup>2</sup> | 2.28 – 0.00mm |
| Dorsolateral striatum (DLS) | 1.0mm<sup>2</sup> | 2.28 – 0.00mm |
| Lateral septum (LS) | 0.4mm<sup>2</sup> | 2.16 – 0.00mm |
| Central amygdala (CeA) | 0.4mm<sup>2</sup> | -1.56 - -2.76mm |
| Basolateral amygdala (BLA) | 0.4mm<sup>2</sup> | -1.80 - -2.76mm |

### Statistical Analysis

Overall social and locomotor activity were assessed using separate for each sex a 2 (Ethanol Social Responding Phenotype: facilitated, inhibited) X 3 (Ethanol Dose: 0, 0.5, 0.75 g/kg) analyses of variance (ANOVAs), with Ethanol Dose treated as a repeated measure. Bonferroni’s multiple comparisons tests were used for further assessment of main effects and interactions. The relationships between overall social activity and neuronal activation indexed via β-gal expression were assessed using linear regressions conducted separately for each ROI within each sex. Effects of Ethanol Social Responding Phenotype on β-gal+ cell counts in each ROI were assessed by independent t-tests separately for males and females. Correlation analyses were used to determine the directionality and strength of the relationship between ROI and behavior. Pearson correlations were then used to evaluate functional connectivity between brain regions for facilitated or inhibited subjects within each sex. A false discovery rate (FDR) of 5% was not implemented given subsequent graph theoretical analysis.

### Graph Theoretical analysis

To perform graph analyses on ethanol social networks, results from Pearson correlations matrices for each pair of brain regions were used to establish centrality measures of degree and betweenness with r coefficients as weights between each pair for facilitated or inhibited subjects within each sex. Degree measures represent the number of edges associated with a node, whereas betweenness represents the shortest pathlength of a node. Only significant correlations were used for subsequent graph analysis using NetworkX, a Python-based graph production script. Girvan-Newman was used to for community detection, and centrality measures were then used to establish hubs regions that potentially mediate interactions between various components of the network. A false discover rate (FDR) was not implemented to incorporate all statistically significant functional connections within clusters.

## Results

### Overall social activity in socially facilitated and socially inhibited by 0.75 g/kg ethanol males and females

A 2 (Ethanol Social Responding Phenotype) X 3 (Ethanol Dose) repeated measures ANOVA of male overall social activity revealed a main effect of Ethanol Dose, (F _2,28_ = 5.075, p < 0.05), and Ethanol Dose by Ethanol Social Responding Phenotype interaction, (F _2,28_ = 23.94, p < 0.0001). As expected, based on a tertile split following administration of the 0.75 g/kg dose, socially facilitated male subjects demonstrated significant increases and socially inhibited males showed significant decreases in overall social activity relative to levels of social activity evident in each phenotype following saline injection (see Fig.2 A), with no effects evident at the 0.5 g/kg dose. Socially facilitated and socially inhibited by ethanol males differed in response to saline, with inhibited by ethanol subjects initially showing significantly greater overall social activity that their socially facilitated counterparts. At the 0.75 g/kg ethanol dose, overall social activity in socially facilitated males was significantly higher than in their socially inhibited counterparts (Figure 2A).

**Figure 2.**
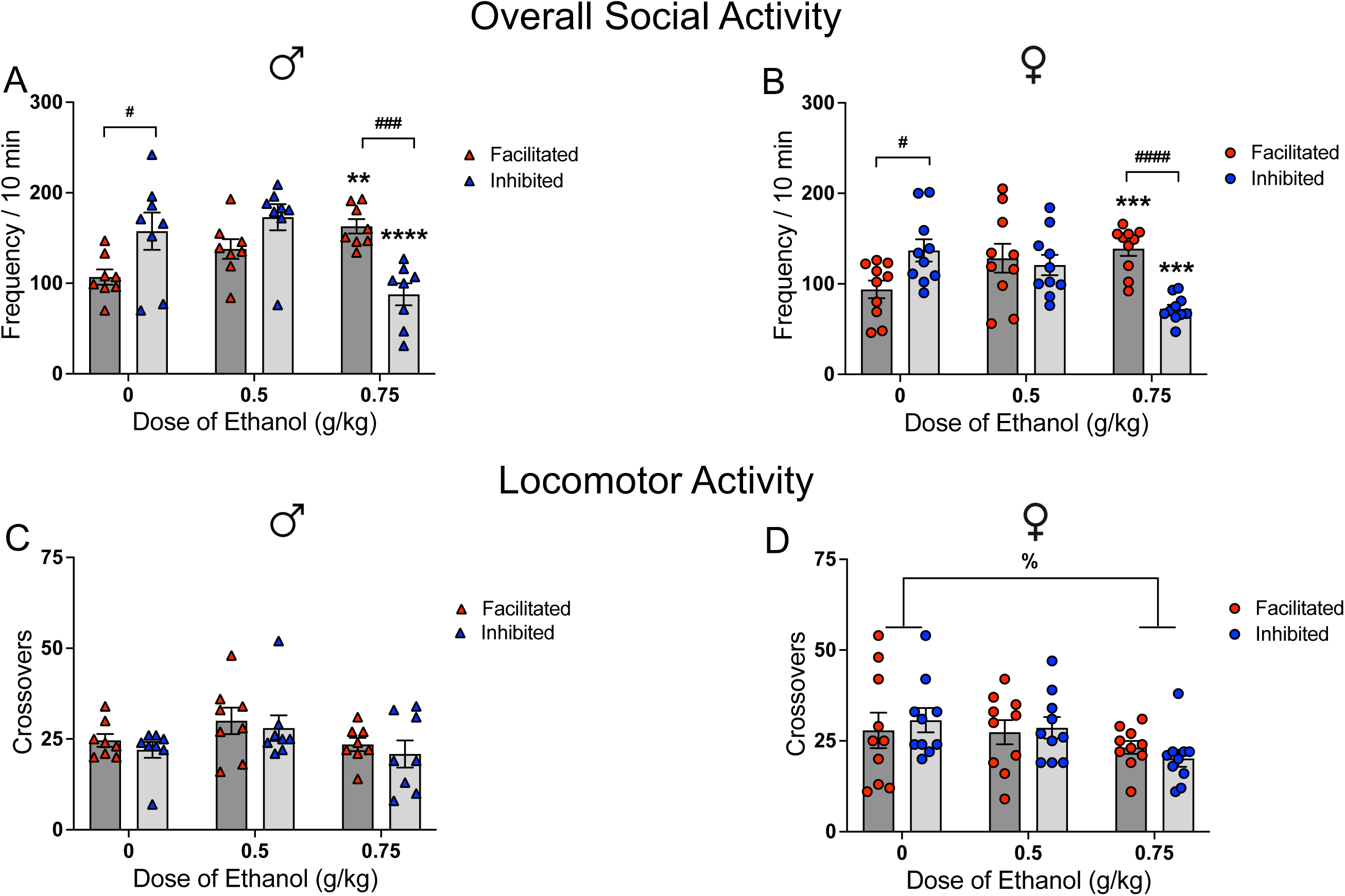
Overall social and locomotor activity in socially facilitated and socially inhibited by 0.75 g/kg ethanol males (A, C) and females (B, D). Significant effects of ethanol relative to saline within each ethanol social responding phenotype are marked with (*): ** - p < 0.01, *** - p < 0.001, **** - p < 0.0001. Significant differences between phenotypes at a certain ethanol dose are marked with (#): # - p < 0.05, ### - p < 0.001, #### - p < 0.0001. Significant effects of ethanol on locomotor activity, with data collapsed across social responding phenotype, are marked with %, p < 0.05.

A 2 (Ethanol Social Responding Phenotype) X 3 (Ethanol Dose) repeated measures ANOVA of female overall social activity also revealed an interaction of Ethanol Social Responding Phenotype and Ethanol Dose, (F _2,36_ = 17.49, p < 0.0001). Socially facilitated females demonstrated significant increases in overall social activity following 0.75 g/kg ethanol, whereas socially inhibited females showed decreases in overall social activity relative to levels seen after saline administration at 0.75 g/kg ethanol. Like males, socially facilitated by ethanol females showed significantly lower social activity than their socially inhibited by ethanol counterparts following saline injection, whereas socially facilitated females had greater levels of social activity than socially inhibited females following 0.75 g/kg ethanol (Figure 2B).

### Locomotor activity in socially facilitated and socially inhibited by 0.75 g/kg ethanol males and females

We next determined whether ethanol-induced social facilitation or social inhibition was associated with ethanol-induced changes of locomotor activity indexed via total number of chamber crossovers. In males, a 2 (Ethanol Social Responding Phenotype) X 3 (Ethanol Dose) repeated measures ANOVA revealed that locomotor activity did not differ as a function of either Ethanol Social Responding Phenotype or Ethanol Dose (Figure 2C). In females, however, a two-way repeated measures ANOVA of locomotor activity revealed a main effect of Ethanol Dose, (F _2,36_ = 3.56, p < 0.05), with 0.75 g/kg ethanol reducing locomotor activity relative to saline (Figure 2D). Taken together, this finding suggests that adolescent females are sensitive to motor impairing effects of ethanol, but that ethanol-induced social facilitation is not related to ethanol effects on locomotor activity.

### Relationship between overall social activity and neuronal activation

We used linear regression and correlation analyses for assessment of relationships between social activity and neuronal activation of the ROIs. In males, overall social activity predicted β-gal expression in the PrL (R^2^ = 0.283, F_1,14_ = 5.53, p < 0.05), IL, (R^2^ = 0.266, F_*-++-+_ = 5.065, p < 0.05), pCL (R^2^ =0.431, F_1,12_ = 9.08, p < 0.05), and NAcSh (R^2^=0.306, F_1,14_ = 6.176, p <0.05). Correlation analyses revealed moderate negative relationships between social activity and neuronal activation of these ROIs (r =-0.532 for PrL, r = - 0.515 for IL, r = - 0.656 for pCL, and r = - 0.553 for NAcSh). No other significant relationships were identified in males (see Table 2). In contrast to social activity, linear regressions revealed no relationships between locomotor activity and β-gal expression in all ROIs (see Table 2). Together, these findings suggest that changes in neuronal activation are primarily associated with social behavior.

**Table 2.** The relationship between behavior and the number of β-gal positive cells for each region of interest assessed with Pearson correlation and linear regression analyses.

| ROI | Social Activity |  |  |  |  |  | Locomotor Activity |  |  |  |  |  |
| --- | --- | --- | --- | --- | --- | --- | --- | --- | --- | --- | --- | --- |
|  | male |  |  | female |  |  | male |  |  | female |  |  |
|  | *r* | *R*<sup>2</sup> | *p* | *r* | *R*<sup>2</sup> | *p* | *r* | *R*<sup>2</sup> | *p* | *r* | *R*<sup>2</sup> | *p* |
| IOFC | - 0.336 | 0.113 | 0.204 | 0.444 | 0.197 | 0.065 | 0.117 | 0.014 | 0.691 | IOFC | 0.027 | 0.515 |
| vOFC | - 0.432 | 0.223 | 0.065 | 0.309 | 0.096 | 0.227 | 0.010 | 0.000 | 0.973 | - 0.132 | 0.017 | 0.615 |
| PrL | **- 0.532** | **0.283** | **0.034** | **0.488** | **0.238** | **0.034** | 0.098 | 0.009 | 0.739 | 0.013 | 0.000 | 0.956 |
| IL | **- 0.515** | **0.266** | **0.041** | **0.476** | **0.227** | **0.034** | 0.046 | 0.002 | 0.875 | 0.250 | 0.063 | 0.287 |
| aCL | - 0.285 | 0.081 | 0.285 | **0.489** | **0.239** | **0.034** | 0.271 | 0.074 | 0.348 | 0.118 | 0.014 | 0.698 |
| pCL | **- 0.656** | **0.431** | **0.011** | 0.434 | 0.188 | 0.056 | - 0.281 | 0.079 | 0.353 | - 0.333 | 0.111 | 0.151 |
| NAcC | - 0.032 | 0.010 | 0.233 | 0.333 | 0.111 | 0.150 | 0.312 | 0.074 | 0.653 | - 0.176 | 0.031 | 0.457 |
| NAcSh | **- 0.553** | **0.306** | **0.026** | **0.563** | **0.317** | **0.015** | - 0.158 | 0.025 | 0.591 | 0.033 | 0.001 | 0.897 |
| DMS | - 0.259 | 0.067 | 0.333 | **0.730** | **0.532** | **0.000** | 0.183 | 0.033 | 0.531 | 0.150 | 0.022 | 0.528 |
| DLS | - 0.077 | 0.006 | 0.775 | **0.852** | **0.726** | **0.000** | 0.304 | 0.092 | 0.291 | 0.076 | 0.006 | 0.749 |
| LS | - 0.248 | 0.061 | 0.355 | **0.555** | **0.308** | **0.013** | 0.154 | 0.024 | 0.600 | **0.464** | **0.216** | **0.045** |
| CeA | - 0.111 | 0.012 | 0.683 | 0.215 | 0.051 | 0.353 | 0.195 | 0.038 | 0.505 | - 0.124 | 0.027 | 0.515 |
| BLA | - 0.251 | 0.063 | 0.348 | 0.423 | 0.179 | 0.071 | 0.243 | 0.059 | 0.402 | 0.044 | 0.002 | 0.857 |

In females, regression analyses revealed that overall social activity predicted neuronal activation of the PrL (R^2^ = 0.238, F_1, 17_ = 5.318, p < 0.05), IL (R^2^ = 0.227, F_1, 18_ = 5.270, p < 0.05), aCL (R^2^ = 0.239, F_1, 17_ = 5.335, p < 0.05), NAcSh (R^2^ = 0.317, F_1, 16_ = 7.417, p < 0.05), DMS (R^2^ = 0.532, F_1, 18_ = 20.50, p < 0.001), DLS (R^2^ = 0.726, F_1, 18_ = 47.77, p < 0.0001), and LS (R^2^ = 0.3080, F1, _17_ = 7.568, p = 0.0136,), with positive correlations evident between overall social activity and β-gal+ cell counts in these ROIs (r = 0.488 for Prl, r = 0.476 for IL, r = 0.489 for aCL, r = 0.563 for NAcSh, r = 0.730 for DMS, r = 0.852 for DLS, and r = 0.555 for LS)). No other significant relationships between social activity and β-gal+ cell counts were evident in females (see *Table 2*). In contrast to males, there was a positive relationship between locomotor activity and β-gal expression in the LS (R^2^ = 0.216, F_1, 17_ = 4.677, r = 0.464, p < 0.05), whereas locomotor activity did not predict neuronal activation of other female ROs (see Table 2).

### Neuronal activation in socially facilitated and socially inhibited males and females

To determine whether neuronal activation of any ROI differed between socially facilitated and inhibited by ethanol subjects, we compared cell counts in multiple brain areas (see Table 3). In males, number of β-gal+ cells in almost all ROIs did not differ between socially facilitated and socially inhibited subjects. In the pCL, however, socially inhibited males had significantly more β-gal+ cells than their socially facilitated counterparts (*t*_12_ = 2.291, *p* < 0.05, Table 3). Conversely, females that were socially inhibited by ethanol generally had lower neuronal activation of the PrL (*t*_17_ = 2.276, *p* < 0.05), IL (*t*_18_ = 2.737, *p* < 0.05), aCL (*t*_17_ = 2.602, *p* < 0.05), NAcSh (*t*_16_ = 2.835, *p* < 0.05), DMS (*t*_18_ = 3.255, *p* < 0.01), DLS (*t*_18_ = 3.666, *p* < 0.01), and LS (*t*_17_ = 2.608, *p* < 0.05) than their socially facilitated counterparts (see Table 3). Number of β-gal+ cells other ROIs did not differ between the two phenotypes. These data support sex differences in regional neuronal activation associated with ethanol-induced social facilitation and social inhibition.

**Table 3.** Number of β-gal+ cells per mm^2^ within each region of interest in socially facilitated and socially inhibited by ethanol adolescent males and females.

| ROI | Male |  |  |  | Female |  |  |  |
| --- | --- | --- | --- | --- | --- | --- | --- | --- |
| | Facilitated<br>Mean ± SEM | Inhibited<br>Mean ± SEM | *t* | *p* | Facilitated<br>Mean ± SEM | Inhibited<br>Mean ± SEM | *t* | *p* |
| IOFC | 210.3 ± 8.9 | 225.9 ± 17.1 | 0.813 | 0.430 | 219.5 ± 14.3 | 187.5 ± 7.6 | 1.830 | 0.086 |
| vOFC | 260.0 ± 14.2 | 304.0 ± 26.8 | 1.454 | 0.168 | 309.4 ± 21.6 | 268.8 ± 11.9 | 1.587 | 0.133 |
| PrL | 148.5 ± 11.3 | 181.6 ± 21.4 | 1.378 | 0.190 | **183.4 ± 15.8** | **143.3 ± 6.0** | **2.276** | **0.036** |
| IL | 128.2 ± 7.2 | 155.3 ± 16.3 | 1.522 | 0.150 | **169.9 ± 11.4** | **133.3 ± 6.9** | **2.737** | **0.013** |
| aCL | 93.6 ± 5.0 | 96.4 ± 8.2 | 0.289 | 0.777 | **100.8 ± 6.8** | **82.3 ± 2.8** | **2.602** | **0.019** |
| pCL | **52.9 ± 3.2** | **64.5 ± 3.9** | **2.291** | **0.040** | 64.2 ± 3.5 | 57.8 ± 3.1 | 1.366 | 0.189 |
| NAcC | 152.7 ± 8.2 | 173.1 ± 20.1 | 0.911 | 0.377 | 181.4 ± 9.1 | 159.1 ± 13.9 | 1.341 | 0.197 |
| NAcSh | 107.8 ± 6.5 | 127.4 ± 12.5 | 1.384 | 0.185 | **131.8 ± 8.0** | **97.8 ± 9.0** | **2.835** | **0.012** |
| DMS | 763.1 ± 45.4 | 880.9 ± 122.9 | 0.899 | 0.384 | **876.7 ± 48.5** | **696.5 ± 26.7** | **3.255** | **0.004** |
| DLS | 640.1 ± 48.8 | 685.7 ± 138.3 | 0.311 | 0.760 | **633.2 ± 40.8** | **468.0 ± 19.1** | **3.666** | **0.002** |
| LS | 103.6 ± 4.7 | 110.3 ± 6.9 | 0.799 | 0.437 | **114.7 ± 5.8** | **97.7 ± 3.4** | **2.608** | **0.018** |
| CeA | 77.8 ± 3.8 | 84.1 ± 12.7 | 0.470 | 0.646 | 84.2 ± 5.2 | 75.1 ± 9.1 | 0.888 | 0.387 |
| BLA | 45.2 ± 2.5 | 47.7 ± 5.7 | 0.397 | 0.697 | 60.3 ± 2.9 | 52.9 ± 5.0 | 1.321 | 0.204 |

Bold text represents significant differences.

### Patterns of neuronal activation: Correlation matrix

We also assessed functional connectivity among ROIs in socially facilitated and socially inhibited by ethanol males and females using Pearson correlation matrices. In socially inhibited males, 70 strong (r between 0.6 and 0.8) and very strong (r > 0.8) positive correlations were evident among almost all cortical and subcortical ROIs (see Fig.3A), with very few correlations not reaching statistical significance. In socially facilitated males, however, only 30 significant positive correlations were evident among ROIs, with the CeA showing no functional connectivity (see Fig.3A).

**Figure 3.**
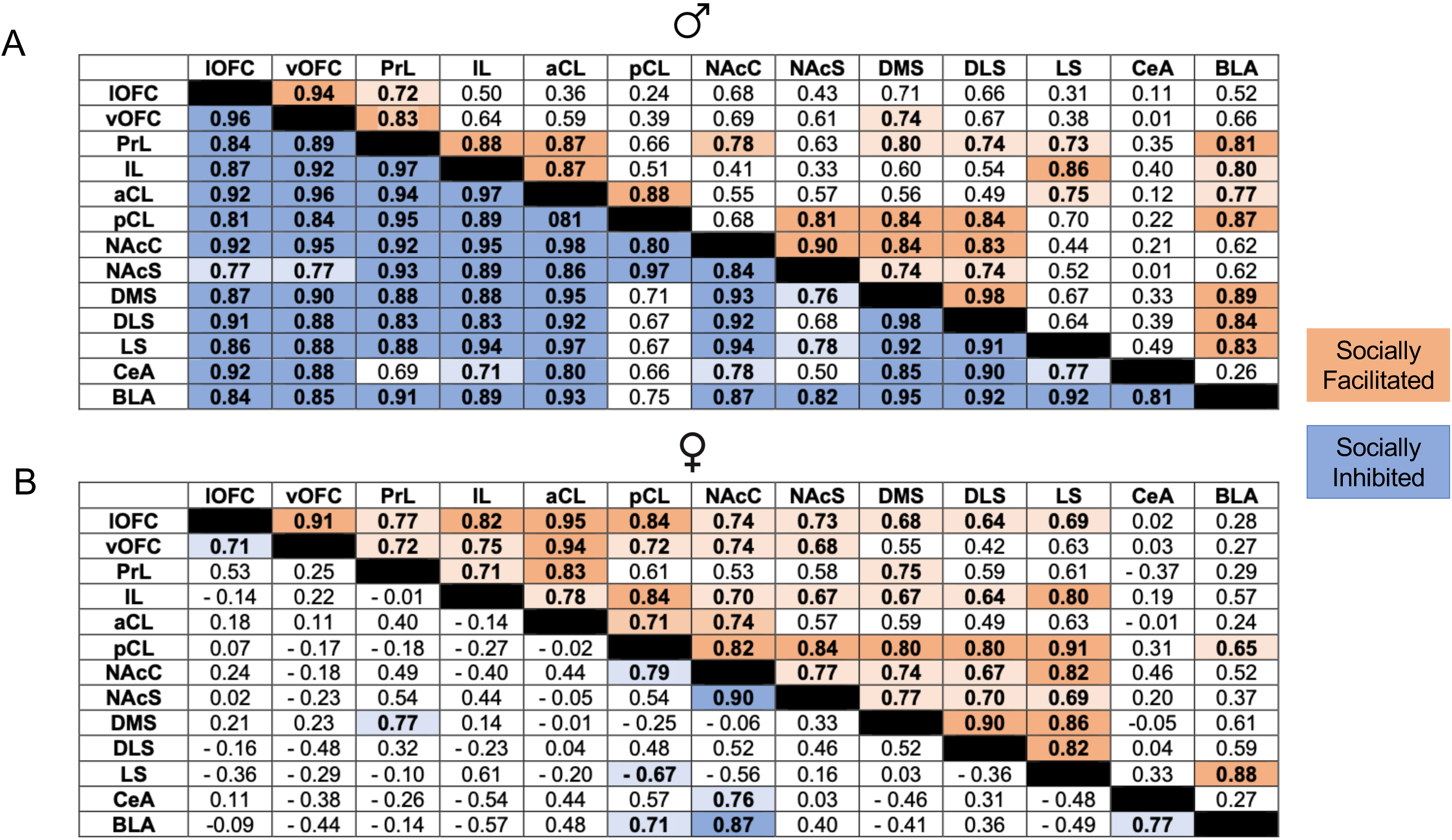
Correlation matrices as a measure of functional connectivity between brain regions in socially facilitated and socially inhibited by ethanol males (A) and females (B). Significant (p < 0.05) strong (r > 0.7) and very strong (r > 0.8) correlations are presented in bold, numbers represent r values.

In contrast to males, only nine significant correlations were evident in socially inhibited females (Fig.3B), with significant positive correlations evident between cortical (vOFC, lOFC, PrL) and subcortical regions (DMS, NAcC, NAcSh,BLA and CeA). A significant negative correlation was also evident between the pCL and LS, with the IL, aCL, and DLS showing no functional connectivity. However, 45 strong and very strong positive correlations that reached statistical significance were evident among ROIs in socially facilitated females, with the CeA showing no functional connectivity (see Fig. 3B).

Finally, we used graph analysis of functional connectivity findings to determine potential hub regions in male and females socially facilitated and socially inhibited by ethanol. Centrality measures pinpointed the PrL as a hub region in males facilitated by ethanol, whereas the pCL was identified as a hub in females (see Fig. 4A and C). Analyses further indicated that the CeA was functionally disconnected in both males and females facilitated by ethanol. In contrast, males inhibited by ethanol did not have a single hub region, but rather five regions that were equally weighted including the NAcC, IL, vOFC, lOFC, and aCL. For females inhibited by ethanol, the NAcC was also identified as a central hub region. Taken together, these findings support a potentially conserved functional disconnect of the CeA in facilitated subjects, and the necessity of the NAcC in inhibited subjects despite previously identified sex differences in regional neuronal activation.

**Figure 4.**
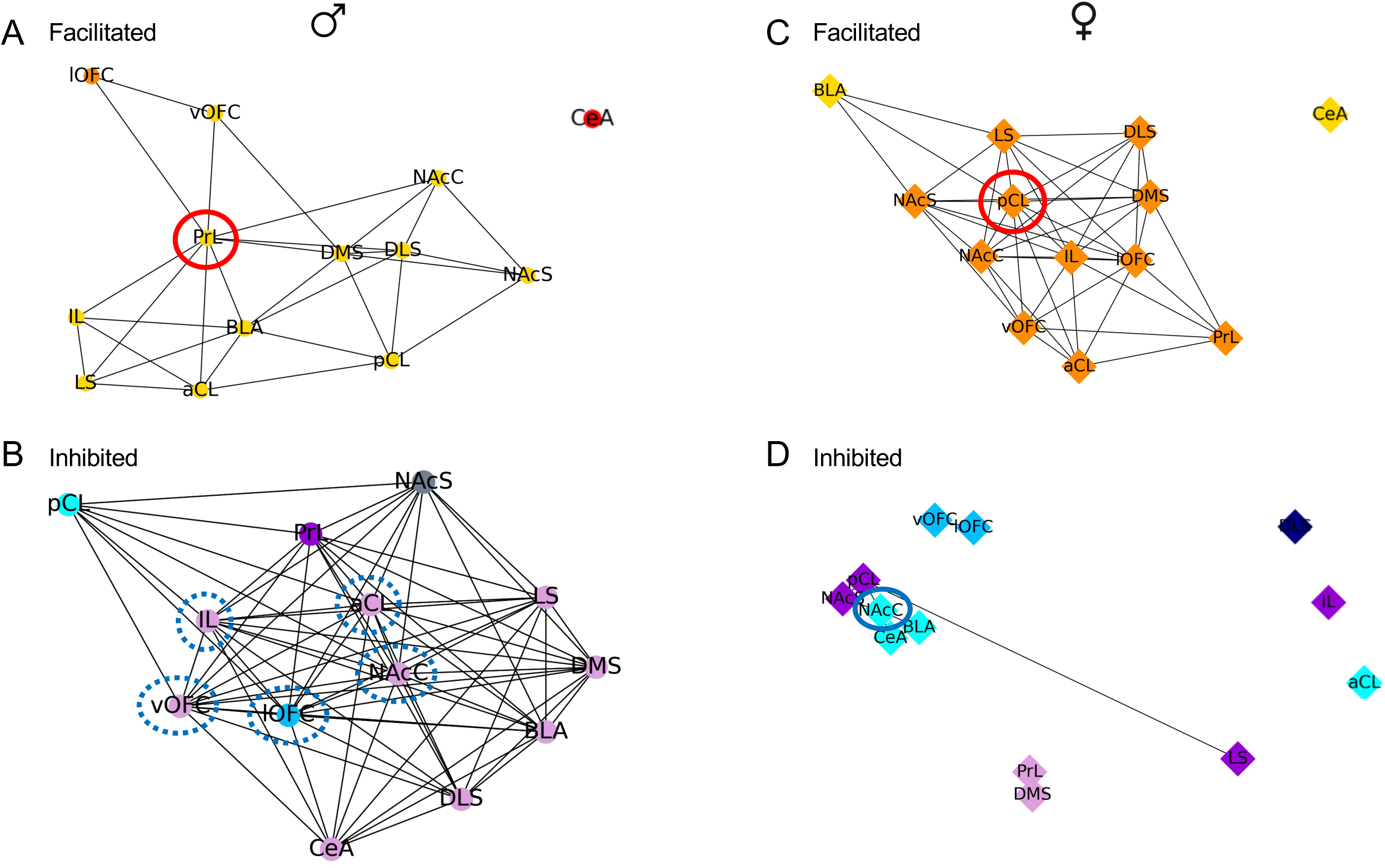
Graph analysis of identified hub brain regions in social facilitated males (A) and females (C) and socially inhibited males (B) and females (D). Identified single hub regions withing a cluster are identified by solid red (facilitated) or blue (inhibited) circles. Dotted circles represented equally-weighted hub regions.

## Discussion

As in our earlier research (Varlinskaya et al. 2015), the present study revealed individual differences in ethanol responsiveness among adolescent cFos-LacZ rats. Behaviorally, socially facilitated and socially inhibited by ethanol subjects differed in response to saline regardless of sex, with inhibited by ethanol subjects initially demonstrating greater overall social activity than their socially facilitated counterparts. This finding suggests that the levels of social responding following acute saline challenge may predict individual sensitivity to socially facilitating and/or socially inhibiting effects of ethanol during adolescence.

In humans, there are substantial individual differences in sensitivity to rewarding, stimulant, and sedative effects of alcohol, with individual alcohol sensitivity playing a rather important role in binge drinking and further AUD development. For instance, in adult heavy drinkers (24 -35 years of age), alcohol has been reported to elicit higher stimulant and rewarding responses and lower sedative responses relative to low drinkers, with greater positive and lower sedative consequences after alcohol consumption predicting high frequency of binge drinking among these heavy drinkers (King et al. 2020). Individuals who experience less aversive effects of alcohol intoxication are more likely to engage in binge drinking and develop AUD later in life (King et al. 2020). Enhanced sensitivity to positive effects has been shown to be a risk factor for developing AUD as well (Morean and Corbin 2010). Animal studies have shown that individual differences in ethanol sensitivity can be investigated in laboratory rodents using ethanol-induced locomotor stimulation, ethanol-induced motor impairment and sedation as well as ethanol-induced taste aversion (Parker et al. 2020). The main disadvantage of these animal models is that animals are often housed and always tested alone, whereas human adolescents predominantly drink with peers. Given that young individuals drink most often in social situations, individual sensitivity to the socially facilitating and/or inhibiting effects of alcohol may be particularly as, if not more important.

Addressing this important issue in the present study, we found that both male and female adolescent rats showed individual differences in sensitivity to the social effects of ethanol. However, these behaviors were accompanied by pronounced sex differences in neuronal activation patterns across the ethanol responding phenotypes. Socially inhibited females had lower neuronal activation of the PrL, IL, aCl, NAcSh, DMS, DLS, and LS than their socially facilitated counterparts, whereas socially inhibited males had more β-gal+ cells in the pCl than those facilitated by ethanol. Furthermore, in males, the analyses of relationships between overall social activity and neuronal activation of the ROIs revealed that social behavior was *negatively correlated* with neuronal activation of the PrL, IL, pCL, and NAcSh (see Table 2), such that lower social activity demonstrated predominantly by socially inhibited by ethanol males corresponded to greater activation of these ROIs. These negative relationships evident in males were rather unexpected, given that activation of the medial prefrontal cortex (mPFC) and nucleus accumbens measured by c-fos expression has been demonstrated in adolescent male rats exposed to a social partner (van Kerkhof et al. 2014). However, in the van Kerkhof et al. study, non-manipulated adolescent males were socially deprived for 24 hours prior to a 15-minute exposure to a social partner, whereas in the present study all experimental subjects were injected with 0.75 g/kg ethanol and socially deprived for only 30 minutes during the habituation period. Indeed, the negative correlations between social behavior and neuronal activation of the mPFC as well as nucleus accumbens may reflect ethanol-induced behavioral and neural changes rather than the involvement of these ROIs in modulation of peer-directed social behavior in adolescent male rats under normal circumstances. In adolescent females, however, the reverse was found relative to males as overall social activity was *positively correlated* with neuronal activation of both the PFC and nucleus accumbens shell, with significantly less β-gal+ cells evident in the PrL, IL, and NAcSh of socially inhibited by ethanol females subjects than in their socially activated counterparts (see Table 3), with lower social activity of socially inhibited by ethanol females corresponding to lower activation of the mPFC and nucleus accumbens. Together, these findings suggest sex-specific involvement of the prefrontal cortical regions and nucleus accumbens in modulation of social behavior in adolescent rats challenged with ethanol.

Responsiveness to rewarding stimuli (including social reward) is related to the mesolimbic system, with the NAc playing a critical role in social reward during adolescence (Trezza et al. 2011; Trezza et al. 2012) and in adulthood (Dolen et al. 2013), with its activity modulated by the mPFC (Del Arco and Mora 2008). Together, results from human and animal studies allow researchers to view the PFC in having a prominent role of the social brain, with prefrontal disfunction resulting in social impairment (Yizhar and Levy 2021). Sex differences in relationships between social activity and activation of certain ROIs, with negative relationships evident in males and positive relationships seen in females, suggest that types of activated cell populations may be different in adolescent males and females challenged with ethanol. It will be important, therefore, to further evaluate the neuronal phenotypes of activated cells within these ROIs.

In females, but not in their male counterparts, positive correlations were also evident between overall social activity and neuronal activation of the medial and lateral dorsal striatum, suggesting the strong relationships (Akoglu 2018). This finding was rather unexpected, although some studies have reported that the dorsal striatum can serve as a neural substrate of social interaction, with functional alterations in the dorsal striatum promoting social deficits in adult mice (Fuccillo 2016; Kim et al. 2017; Peca et al. 2011). Importantly, in a study that included both male and female mice, depletion of both parvalbumin-expressing fast-spiking interneurons and large cholinergic interneurons in the dorsal striatum produced marked social deficits in male mice, but not their female counterparts (Rapanelli et al. 2017), suggesting sex-specific roles of these interneurons in social interaction. The present study is the first to find sex-specific involvement of the dorsal striatum in modulation of social responding in adolescent females acutely challenged with ethanol, with greater neuronal activation of the DLS and DMS corresponding to higher social activity demonstrated by socially facilitated by ethanol females. Given that subtypes of activated neurons cannot be identified with X-gal staining, the next step of our research is to identify subtypes of activated neurons in the dorsal striatum of socially facilitated and socially inhibited by ethanol adolescent rats.

In adolescent females challenged with ethanol, overall social activity was correlated positively with activation of the aCL as well as of the LS, although these significant correlations were not as strong as between social activity and neuronal activation of the DMS and DLS. In males, however, overall social activity was negatively correlated with activation of the pCL. The claustrum (CL) has extensive connections with multiple cortical and subcortical brain regions, including the PFC and amygdala (Jackson et al. 2018; Smith et al. 2020). Although possible CL involvement in social behavior modulation as well as in ethanol effects has not been studied in laboratory rodents, a subpopulation of CL neurons has been shown to modulate stress-induced anxiety-like behavior of male mice through interaction with the BLA (Niu et al. 2022; Tanuma et al. 2022). However, an important issue of CL involvement in regulation of social interaction and/or anxiety in adolescent rats of both sexes under basal conditions as well as following ethanol challenge still needs to be addressed in future experiments. In contrast, extensive studies of the LS suggest its fundamental role in regulating social interaction (reviewed in (Menon et al. 2022), with the LS also being among brain regions showing consistent ethanol-induced activation (Vilpoux et al. 2009). Therefore, the relationship between neuronal activation of the LS and overall social activity in adolescent females challenged with ethanol was not surprising. However, the lack of such a relationship in males was rather unexpected. The LS was also the only ROI in which neuronal activation was positively correlated with locomotor activity (number of movements between the compartments) in adolescent females, suggesting sex-dependent involvement of the LS in ethanol-induced changes of social locomotor activity during adolescence. Importantly, all other ROIs did not correlate with locomotor activity further supporting their involvement in the social brain network.

Differences between ethanol responding phenotypes as well as sex differences became even more apparent when the correlation analysis was used for further assessments of neuronal activation patterns. In socially inhibited males, strong correlations were evident among almost all ROIs (90%), suggesting synchronized activation of these regions, whereas markedly fewer correlations among ROIs (38%) were seen in socially facilitated by ethanol adolescent males. In males, overall social activity was negatively correlated with neuronal activation of the PrL, IL, pCL, and NAcSh (see Table 2), suggesting that activation of these brain regions may be involved in ethanol-induced social suppression. Indeed, very strong correlations (r values between 0.89 and 0.97) were evident among these ROIs in socially inhibited by ethanol adolescent males. In socially facilitated males, however, strong correlations were seen between the PrL and IL as well as between aCL and NAcC, while activation of the prelimbic and infralimbic parts of the mPFC was not correlated with activation of the pLC and NAcSh. Furthermore, in socially inhibited by ethanol males, strong correlations were evident between the orbitofrontal cortex (lOFC, vOFC) and amygdala (CeA, BLA), with no such correlations seen in socially facilitated by ethanol adolescent males. To the extent that the observed correlations can be viewed as an index of functional connectivity (Cross and Leslie 2021; Tanimizu et al. 2017), this finding may have an important translational value. In human adolescents, a relationship between amygdala-orbitofrontal connectivity and past month alcohol use has been reported, with lower connectivity associated with higher alcohol consumption among males (Peters et al. 2015).

In contrast to socially inhibited males, their female counterparts had very few significant correlations among ROIs (10%, see Figure 3), with positive correlations evident within the OFC, nucleus accumbens, and amygdala. Less sex differences were evident in patterns of neuronal activation in subjects socially facilitated by ethanol. In general, socially facilitated females showed more functional connectivity (58% vs 38%) than their male counterparts. In socially facilitated by ethanol females, strong correlations were seen between neuronal activation of certain cortical regions (lOFC, IL, but not PrL) and all other ROIs excluding the CeA and BLA. In socially facilitated males, however, neuronal activation of the PrL was correlated with activation of multiple ROIs except for the pCL, NAcSh, and CeA. Furthermore, while in socially facilitated females, neuronal activation of the LS was correlated with activation of multiple ROIs, whereas in males, the LS showed functional connectivity only with the PrL, IL, and aCL. The BLA of socially facilitated females showed rather poor functional connectivity, with significant correlations evident with neuronal activation of the pCL and LS. In their male counterparts, however, neuronal activation of the BLA was strongly correlated with activation of the PrL, IL, CL, DMS, DLS, and LS. The observed sex differences in patterns of neuronal activation may reflect differential gating of the social brain circuit in response to ethanol challenge as well as to a social partner in adolescent males and females.

There were also some similarities in male and female patterns of neuronal activation. For instance, neuronal activation of the BLA was not correlated with activation of the CeA in socially facilitated animals of both sexes, whereas this correlation was evident in socially inhibited males and females, suggesting that a synchronized activation of the BLA and CeA might play a role in social suppression and/or social anxiety-like responding regardless of sex. The BLA and CeA are differentially involved in modulation of social behavior. For instance, activation of the BLA suppressed social behavior in primates, whereas inhibition of either the BLA or CeA enhanced social behavior, with this enhancement more evident after inhibition of the BLA (Wellman et al. 2016). Furthermore, neuronal activation of the BLA and NAcC were strongly correlated in socially inhibited males and females, whereas no such correlations were evident in socially facilitated subjects regardless of sex. These findings agree with the results of optogenetic activation of the BLA-NAc circuit in mice, with this activation suppressing social interaction and increasing social avoidance (Folkes et al. 2020). Furthermore, the CeA was the only ROI that showed no functional connectivity in socially facilitated males and females, suggesting very little involvement, if any, of the CeA in ethanol-induced social facilitation. This dissociation is strongly evident in our hub analysis. Given that the CeA plays an important role in physiological and behavioral responsiveness to anxiogenic stimuli (Gilpin et al. 2015), this finding was rather predictable. Interestingly, for socially inhibited by ethanol subjects, the NAcC was determined to be a hub region for both sexes, further supporting the potential role of the NAcC to BLA pathway in social avoidance.

In conclusion, the data presented clearly demonstrate individual and sex-related differences in responsiveness to acute ethanol in adolescent cFos-LacZ rats, but also a convergence on the role of ROIs for subjects socially facilitated or inhibited by ethanol. Neuronal activation patterns associated with ethanol-induced social facilitation and ethanol-induced social inhibition following an acute challenge with 0.75 g/kg ethanol differed in males and females, with neuronal activation of the certain ROIs contributing to social facilitation in females and social inhibition in males. Despite these sex differences, findings also support the dissociation of CeA in facilitated, but integrated involvement of the NAcC in facilitated by ethanol subjects.

## Acknowledgements

Supported by NIH grants P50AA017823, T32 AA025606 (Development and Neuroadaptation in Alcohol and Addictions, DNAA) and F31 AA029300. Any opinions, findings, and conclusions or recommendations expressed in this material are those of the author(s) and do not necessarily reflect the views of the above stated funding agencies. The authors have no conflicts of interest to declare.

## Notes

### Competing Interest Statement

The authors have declared no competing interest.

